# A Quantitative Evaluation of Topological Motifs and Their Coupling in Gene Circuit State Distributions

**DOI:** 10.1101/2022.07.19.500691

**Authors:** Benjamin Clauss, Mingyang Lu

## Abstract

One of the major challenges in biology is to understand how gene interactions collaborate to determine overall functions of biological systems. Here, we present a new computational framework that enables systematic, high-throughput, and quantitative evaluation of how small transcriptional regulatory circuit motifs, and their coupling, contribute to functions of a dynamical biological system. We illustrate how this approach can be applied to identify four-node gene circuits, circuit motifs, and motif coupling responsible for various gene expression state distributions, including those derived from single-cell RNA sequencing data. We also identify seven major classes of four-node circuits from clustering analysis of state distributions. The method is applied to establish phenomenological models of gene circuits driving human neuron differentiation, revealing important biologically relevant regulatory interactions. Our study will shed light on a better understanding of gene regulatory mechanisms in creating and maintaining cellular states.

## Introduction

One of the main questions in systems biology is to understand how complex gene regulatory networks perform their functions to control important biological processes, such as cell differentiation and cell division^1, 2^. Over the years, researchers have focused on studying gene circuit motifs, defined as reoccurring small circuit topologies within larger biological gene regulatory networks^3^. It has been shown, by approaches in synthetic biology^4^, computational systems-biology modeling^5, 6^, and experimental systems biology^7^, that different gene circuit motifs exhibit distinct functions in creating and maintaining circuit states, driving state transitions, and processing signals. For example, an autoregulatory negative feedback loop is known to suppress gene expression noise^8, 9^; a two-node toggle switch circuit can generate bistability^10, 11^; and an incoherent feed-forward loop can achieve adaptation^12^. Although the dynamical behaviors of individual circuit motifs have been widely studied, it is still challenging to characterize the roles of the circuit motifs when they interact with other motifs, or when they present within a large biological network. Due to the presence of additional gene-gene interactions, circuit motifs may behave differently from the standalone motifs. Understanding emergent behaviors arising from motif coupling will greatly improve our understanding of functionality of circuit motifs and lager networks in general.

Gene circuit motifs were classically identified by searching the topology of a large biological network, such those from E. coli and yeast, for the presence of smaller circuit motifs^2, 3, 13^. Motifs are important when they are over-represented in biological networks compared to similarly generated random networks. This approach usually only considers the frequency of circuit motifs’ appearance, but not their functionality, for initial identification. To address this issue, recent studies^5, 6, 14, 15^ have been focused on identifying functionally relevant circuit motifs capable of producing specific dynamical behaviors using mathematical modeling and then analyzing them for enriched motifs. These types of approaches have been devised and applied to elucidate circuits capable of generating oscillations^16,17,18^ and multiple stable steady states^6, 19, 20^. Ye et al.^6^ identified three-node circuits capable of generating stepwise transitions between four states with limited reversibility. Analysis of these circuits allowed them to identify regulatory interactions controlling the development of T-lymphocytes^6^. Schaerli et al.^14^ investigated circuits capable of stripe formation, identifying incoherent feed-forward loops and a two-node motif containing activation and inhibition as the critical motifs. However, there are a still a few questions remain to be addressed for a more general applicability. First, mathematical modeling of gene circuits is often performed with a set of fixed kinetic parameters or examined with parameters sampled from a narrow range, limiting the robustness and accuracy of modeling methods in evaluating circuit behaviors. Second, there lacks a quantitative scoring method allowing the ranking of circuits for *any* desired functionality, or to measure *functional* similarities and differences between two circuits. Third, it is still challenging to evaluate motif coupling, *i.e.*, how one circuit motif interacts with another to produce the desired behaviors. The coupling of circuit motifs has been shown to play important roles to the overall behavior of gene circuits^21, 22^. In particular, the role of circuit coupling may depend on the proportion of shared nodes between the two coupled circuit motifs^5^. To the best of our knowledge, no systematic quantitative analysis is available to statistically evaluate circuit coupling.

To overcome these challenges, we devised a computational framework that allows robust discovery of causal gene circuit motifs and patterns of motif coupling by defining a quantitative score to identify circuits capable of achieving specific functions. Circuit functions can be anything related to the circuit dynamics or steady state distributions, e.g., gene expression allowing three state clusters, specific multivariant distribution of gene expression, and gene expression distributions derived from experimental single cell data. In this study, we performed the first-ever comprehensive analysis on all non-redundant four-node transcriptional regulatory circuits. Compared to previous studies on three-node circuits^6, 14, 23^, our analysis has the following advantages. First, there are around sixty thousand non-redundant four-node circuits (see Methods), which is still manageable to perform extensive computational simulations and is sufficiently large for a robust statistical analysis. Second, analyzing these four-node circuits allows for the evaluation of the roles of individual two-node circuit motifs in larger circuits. Third, analyzing four-node circuits has a major advantage in evaluating the role of the coupling between two two-node circuit motifs. Having four nodes in the larger circuit, we can statistically evaluate whether two two-node motifs are likely to occur in a four-node circuit with or without sharing the same node. This is infeasible from the typical analysis of three-node circuits.

To model the dynamical behaviors of all these circuits in a high throughput way, we applied our recently developed method, random circuit perturbation (RACIPE)^24^, to simulate an ensemble of ODE models with randomly generated kinetic parameters and analyze the steady-state gene expression distribution from these models. RACIPE has been applied to elucidate the dynamics of synthetic gene circuits^10, 21, 25^, gene networks regulating stem cell differentiation^26^, cell cycle^27^, B-cell development^28^, and epithelial-mesenchymal transitions^29, 30^ (EMT). These previous studies have shown that, despite having randomly sampled kinetic parameters and initial conditions, steady state solutions of models generally converge to distinct clusters of gene expression patterns representing the functional states of the circuit. Functional states to which most models converge represent state distributions of the circuit and define its overall behavior. Furthermore, these studies also show that topology has an instructive role in defining the state distribution. We have also previously shown that RACIPE-simulated data resembles single cell gene expression data, yet another advantage for discovering biologically relevant circuits.

In the following, we first give an overview of the computational framework. We then illustrate the methodology with two applications to identify all four-node gene circuits allowing a triangular three-state distribution and a linear three-state distribution. From an enrichment analysis, we can identify the enriched two-node circuit motifs and the patterns of their coupling. Next, we show how this framework can be applied to identify 1) clusters of circuits with distinct gene expression functions and 2) circuits with similar state distributions to any other starting circuit. Finally, we show how our method can be applied to identify circuits motifs and their coupling responsible for experimentally observed single-cell gene expression state distributions.

## Results

### A quantitative method to identify circuits with defined functionalities

In this study, we devised a computational framework that enables us to quantitatively evaluate the functionality of transcriptional gene circuit motifs. We used statistical analysis of large ensembles of simulation data to identify circuits best able to perform specific functions, and then analyze those circuits to identify the associated functional units. A schematic overview of the framework is illustrated in **Fig. 1**. First, we systematically generated all possible four-node gene circuits (**Fig.1A**) (see Methods for details of circuit generation) containing regulatory interactions of transcriptional activation/inhibition between nodes and autoregulation of individual nodes. Only circuits containing four functionally connected nodes were considered for analysis, excluding circuits equivalent to three or less nodes. Moreover, for redundant circuits, *i.e.*, circuits with the same topology but switched gene names, only one was included. Second, to each circuit, we applied RACIPE^24^ to generate an ensemble of 10,000 ODE models with randomly generated kinetic parameters (see Methods for details of circuit simulation). Here, for a node that is transcriptionally regulated by multiple nodes, we assume that the effects of the transcriptional regulations from these nodes are independent, resembling AND logic. From the ensemble of mathematical models, we then evaluated the distribution of the steady-state gene expression (**Fig.1B**). Such a state distribution can be interpreted as analogous to single-cell gene expression distributions driven by the specific gene circuit, incorporating the presence of cell-to-cell variability through the sampling of random kinetic parameters^24^. Different circuit topologies can often be associated with a variety of state distributions depending on the range of kinetic parameters explored, highlighting the need to explore a broad parameter space to better characterize the behavior of a circuit. Third, the core of our approach is to perform statistical analysis on the four-node circuits with similar state distributions (**Fig.1C**). The circuit analysis allows the identification of enriched circuit motifs that are functionally associated with state distributions. We also extended the circuit analysis to identify patterns of coupling between two circuit motifs. We mainly focused on circuit motifs of two nodes, but this approach can be readily extended to analyze circuit motifs of other sizes.

**Figure 1.**
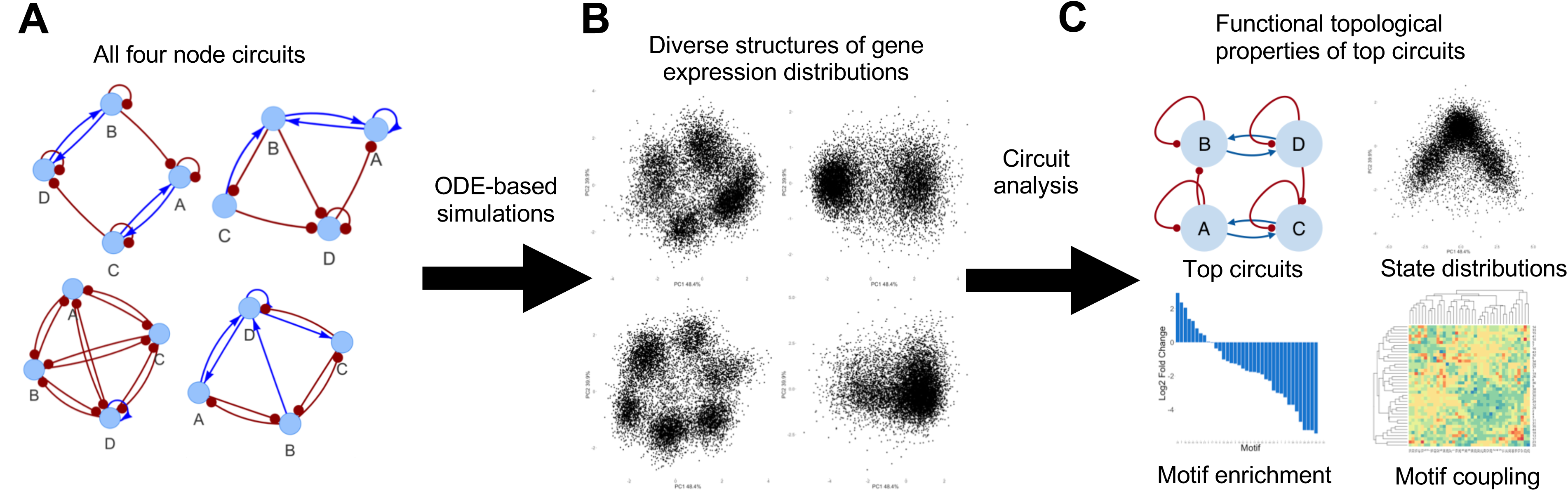
A schematic overview of circuit motif analysis. **(A)** All non-redundant four-node gene circuits are first generated. **(B)** The dynamical behavior of these circuits were then explored using ensemble-based ODE simulations, resulting in diverse structures of state distributions. **(C)** This rich simulation dataset allows us to (1) identify circuits with a certain structure of state distribution, (2) identify reoccurring two-node circuit motifs and their coupling among these circuits, (3) quantify the similarity of circuit functions from state distributions.

### Characterizing Circuits of three states with a triangular or linear state distribution

To illustrate the application of our circuit motif analysis framework, we evaluated four-node circuit topologies capable of generating a triangular arrangement of three gene expression states, as illustrated in **Fig.2A**. This type of triangular state distributions is frequently observed in biological processes involving distinct cellular state transitions of multiple steps, *e.g.*, multi- lineage differentiation from a progenitor cell type to two distinct differentiated cell types, as is frequently observed in hematopoietic lineages ^31, 32^. For each four-node circuit, we applied k-means clustering (k = 3) to the RACIPE simulated gene expression profiles of all the non-redundant circuits and calculated the triangularity score *Q*_1_, as defined in Equation (1) in the Method section. Higher *Q*_1_ values indicate state distributions with a greater degree of separation between the three clusters. We then ranked all non-redundant four-node circuits from high to low *Q*_1_ values, with the top five ranked circuits illustrated in **Fig.2B**. As demonstrated in the PCA projections of the simulated gene expression data of the corresponding circuits (**Fig.2B**, bottom row), these circuits create gene expression state distributions of three states arranged in a triangular shape. Interestingly, the topologies of top ranked circuits are remarkably similar with clear patterns of two-node circuit motifs, such as motif 25 (**Fig.S1** for the list of motifs and their indices) appearing twice in each network without sharing a node and motif 25 and 16 co-occurring while always sharing a node (see below for details).

**Figure 2.**
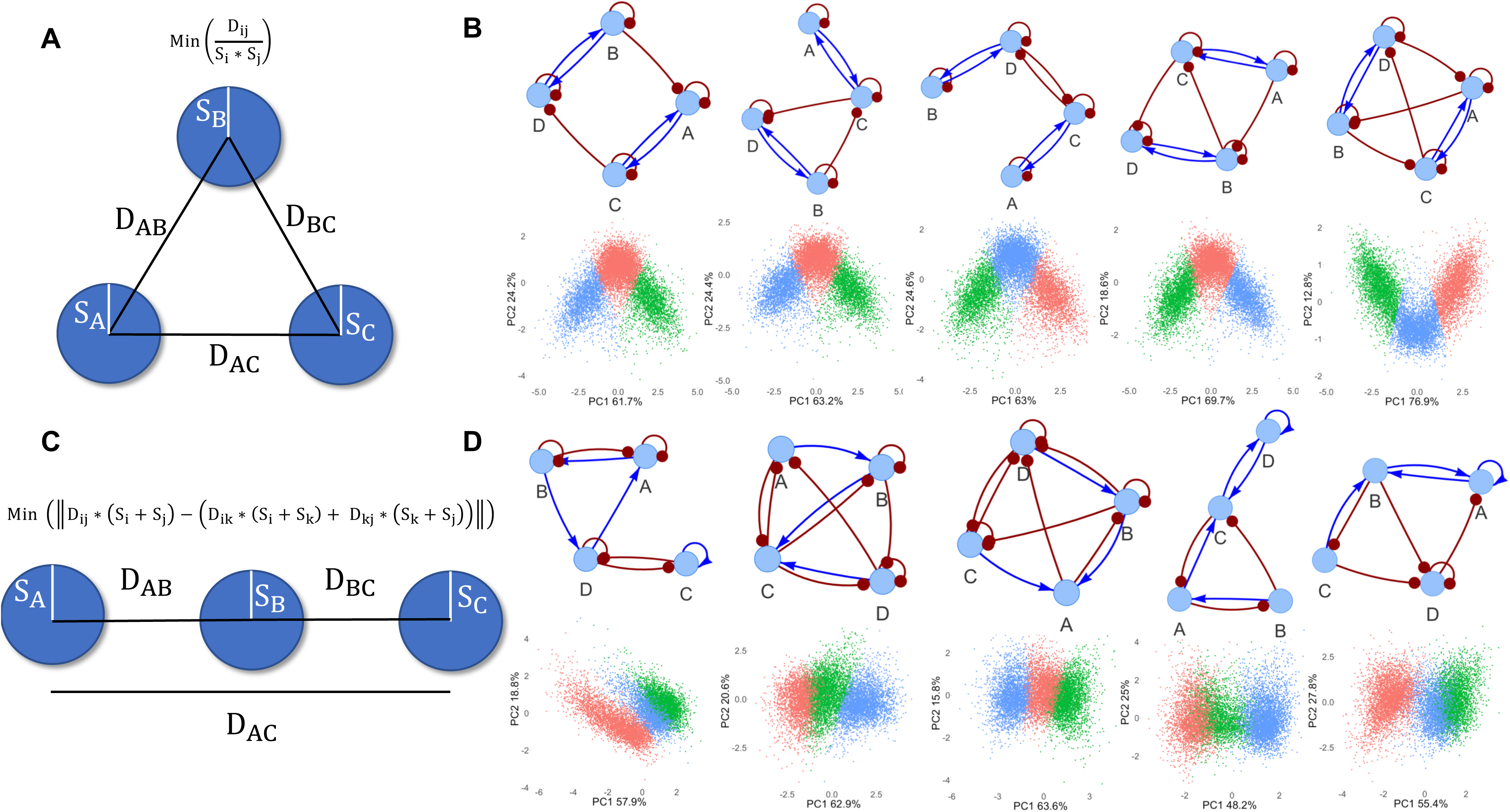
Identifying four-node circuits with triangular and linear state distributions. **(A)** Illustration of the score defined to identify circuits with a triangular state distribution. **(B)** Illustration of the top five circuits with the highest triangularity scores. The plot shows the circuit diagrams (top row) and the scatter plots of the projection of RACIPE simulated gene expression onto the first two principal components (bottom row). In the circuit diagrams, the lines and arrows in blue represent activating interactions; the lines and dots in red represent inhibiting interactions. In the scatter plots, red, blue, and green colors show the three clusters of gene expression states identified by k-means clustering (k=3). **(C)** Illustration of the score defined to identify circuits with a linear state distribution. **(D)** Illustration of the top five circuits with the highest linearity scores.

Next, we explored four-node circuit topologies capable of generating three gene expression states arranged into a linear shape, as illustrated in **Fig.2C**. This type of linear state distributions is frequently observed in the biological processes involving cellular state transitions through an intermediate state, such as transdifferentiation along a singular lineage during Epithelial-mesenchymal transition^33–35^. We performed the same k-means clustering analysis to the RACIPE simulated data, as described above, ranking all non-redundant four-node circuits from low to high *Q*_2_ value, where *Q*_2_ is a linearity score defined in Equation (2, Methods). The top five ranked circuits of linear state distribution are illustrated in **Fig.2D**. We observed that these circuits can indeed produce a linear distribution of three gene expression clusters. The structures of the circuit topologies are similar among them but distinct from those allowing for a triangular state distribution. We observed repeated motifs containing activating and inhibiting edges between the two nodes, in stark contrast to the motifs observed for the triangular score.

Taken together, our results demonstrates that the two scores, *Q*_1_ and *Q*_2_, are effective at detecting circuits capable of producing three states of triangular or linear state distributions. While we have shown an application of this method to the linear and triangular structures, this analysis can be extended to any scoring function defining a particular state distribution, allowing a similar ranking analysis to identify circuits with novel features.

### Enrichment analysis identifies circuit motifs and motif coupling

Next, we evaluated the properties of the circuit topology for those with a triangular state distribution. We first enumerated all possible two-node circuit motifs (see **Fig.S1**) and identified their occurrence in each four-node circuit. We evaluated the enrichment of each two-node motif in circuits with top triangularity scores (about top 1% circuits, see Methods for details), as shown in **Fig.3A**. The topmost enriched circuit motif for the triangular state distribution is a circuit of two genes with both mutual activation and self-inhibition (motif 25). Interestingly, the top three enriched motifs all contain self-inhibition on both genes, suggesting the importance of the inhibitory autoregulation in generating three well separated states. Furthermore, the bottommost enriched circuit motifs are very different from the topmost motifs, in that the motifs are likely to contain activating autoregulation and inhibition between nodes. Interestingly, the most under-enriched motif, motif 21, is similar to motif 25 except that both nodes contain self-activation. This again underscores the importance of negative autoregulation in producing the triangular three state distribution.

**Figure 3.**
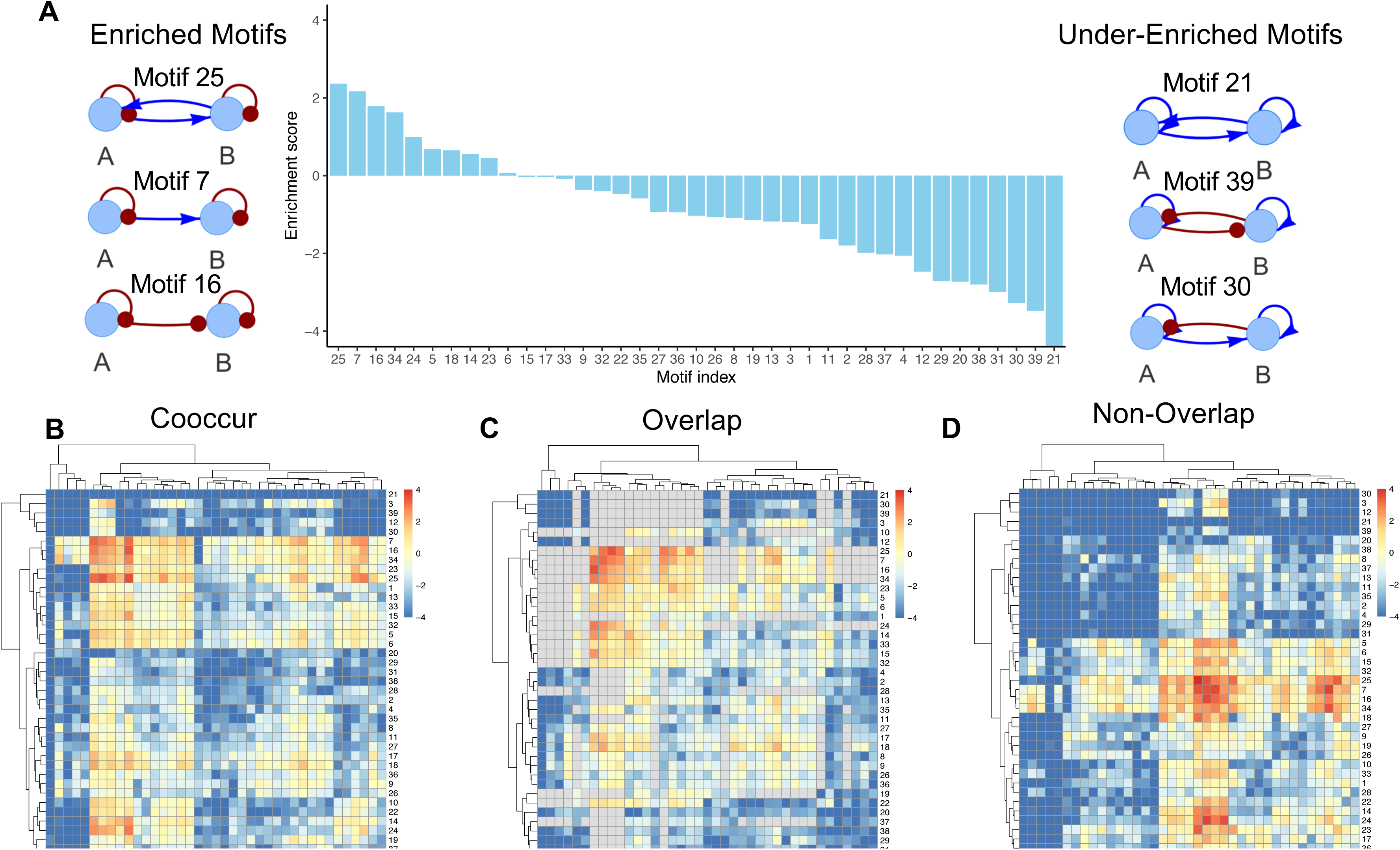
Motif enrichment analysis using the triangularity score. **(A)** The enrichment score for all two-node circuit motifs using the triangularity score. All enrichment results are significant (p-value < 0.05) except for motifs 6, 15, 17, and 33. Panels **(B - D)** show the heatmaps of the enrichment scores for the coupling between two types of two-node circuit motifs. The hierarchical clustering analysis was performed using Euclidean distance and complete linkage method. Panel **(B)** shows the outcomes for all two co-occurring motifs; panel **(C)** for two co-occurring motifs that share the same node; panel **(D)** for two co-occurring motifs that do not share the same node.

To understand how circuit motifs cooperate to generate triangular state distributions, we performed a similar enrichment analysis on the co-occurrence of two motifs among the same top ranked circuits. We can visualize the pattens of circuit coupling from the heatmap of enrichment scores for the co-occurrence of two motifs (**Fig.3B**), for co-occurrence of two motifs with a shared node (motifs with overlapping, **Fig.3C**), and for co-occurrence of two motifs without any overlapping node (**Fig.3D**). Interestingly, the topmost enriched motif coupling patterns are (1) between two motifs of #25 and (2) between motifs 25 and 16. Furthermore, coupling between two motifs of #25 is the highest enriched motif pair for non-overlapping. This is consistent with what we observed in the top four-node circuits with the triangular state distribution – in the top 20 circuits there are 18 cases that contain exactly two motifs of #25 without a shared node (**Fig.S2)**. In addition, we observed that motifs 14, 17, 18, and 24 all have relatively higher enrichment for coupling with motifs 7 and 25 (see **Fig.3B**), despite having relatively lower enrichment of as standalone motif (**Fig.3A**). For motif coupling without a shared node (**Fig.3D**), we observed surprisingly high enrichment between motifs 24 and 34, as well as between motifs 23 and 25. The relatively high enrichment of motifs 14, 17, 18, 23, 24, and 34 in the motif coupling indicates emergent behaviors for these motifs that contribute to the triangular state distribution only when coupled with other specific motifs.

We examined the properties of circuits capable of generating linear state distributions in a similar way to the analysis for the triangular state distribution. Enrichment of motifs in the top 1% of circuits ranked by the linear score were identified, as shown in **Fig.S3**. The topmost enriched motif for the linear score, motif 34, is characterized by two nodes with negative autoregulation and one excitatory and one inhibitory edge between nodes (see **Fig.S1**). The second top motif, motif 33, is similar to the topmost, however only one node contains negative autoregulation.

When compared to the bottom three enriched motifs, once again we observed a striking difference in the nature of auto-regulation; while the top motifs tend to contain negative autoregulation, the bottom motifs contain positive autoregulation. This may point to a general importance for negative autoregulation in the ability to create three state distributions. Indeed, it has been shown that negative autoregulation decreases gene expression noise^3^, which may contribute to the ability of the identified motifs to generate separate states. We also noted that motif 39, a classic example of the self-activating toggle switch circuit capable of generating tristability^10, 11, 36, 37^, is identified as one of the bottom enriched motifs. We believe motif 39 is under enriched because our linear state distributions appear more continuous than those generated by toggle switch with self-activation alone. These findings demonstrate that our method can detect quantitative differences in state distributions that are qualitatively similar (*i.e.*, continuous three state vs disparate three states) and therefore identify more specifically enriched motifs. The coupling of circuit motifs observed in four-node circuits favoring the linear score was shown in **Fig. S2D-F**.

Taken together, our data indicates that we can identify the quantitative contribution of key regulatory interactions, motifs and their coupling, responsible for producing specific structures of gene expression data (**Fig 3**, **Fig.S3**). We show how this can be applied to both linear and triangular arrangements of gene expression states; however, this approach can be expanded to any theoretical state distribution and identify motifs of other sizes.

### Circuit motifs of linear and triangular state distributions are frequently observed in biological networks

Next, we searched for the occurrence of the top enriched motifs in both the linear and triangular scores in PluriNetWork^38^, a manually curated literature-based databases of transcription factor regulations for mouse pluripotency. We identified a total of 57 motifs from both the triangular and linear scores in the pluripotency gene regulatory network – 21 cases of motif 5, 17 of motif 6, seven of motif 14, ten of motif 15, 1 of motif 32, and 1 of motif 25. Among these motifs, Sall4 and Oct4^39^ form motif 25 -- a circuit of two genes with both mutual activations and self-inhibitions, as also supported by regulatory interactions from the TRRUST database^40^. Oct4 is a well-known pan-pluripotency transcription factor, and Sall4 is important in ESC proliferation and differentiation^39, 41, 42^. Tcf7 and Oct4 also form motif 23, a circuit of mutual activation with one node containing positive auto-regulation. Tcf7 is an important transcription factor that alone can restore trilineage differentiation abilities in mouse ESCs lacking all full length TCF/LEFs, demonstrating its importance in generating three states^43^. Note that the PluriNetWork database may have many missing interactions (*e.g.*, only two genes have negative autoregulation) or annotations (*e.g.*, interactions labeled as unknown), therefore the occurrence of circuit motifs are likely underestimated. It is worth further investigating the roles of these circuit motifs in stem cell differentiation.

### Classifying types of state distributions of all four-node circuits

In the previous sections, we relied on defining a score to quantify a particular structure of state distribution and then ranking circuits with the score. Here, we aimed to classify gene circuits by structures of state distributions with an unsupervised top-down approach. To achieve this, we defined a distance function to quantify the differences in state distributions between two four-node circuits, and then applied clustering analysis on the resulting distance matrix. The distance function we chose was based on a multivariate Kolmogorov-Smirnov (KS) statistic, which allows the quantification of the differences between the gene expression distributions of two circuits (details in Methods). We then devised a subsampling approach of Louvain clustering (schematic overview in **Fig. 4A**, details in Methods) for the distances between all non-redundant four-node circuits, from which we identified 20 circuit clusters with more than 10 members representing distinct classes of state distributions. For example, **Fig. 4B** shows representative state distribution from three different clusters – one allowing a single gene expression state (leftmost), one allowing two separated gene expression states (middle), and another allowing a circular state distribution (rightmost). From the histogram of the number of circuits in each cluster (**Fig.4C**), we observe seven major circuit clusters. Interestingly, circuit clusters with more members tend to have simpler state distributions (*i.e.*, distributions with one or two gene expression clusters), while clusters containing more complex structures (*e.g.*, those with six gene expression clusters) often contain less members.

**Figure 4.**
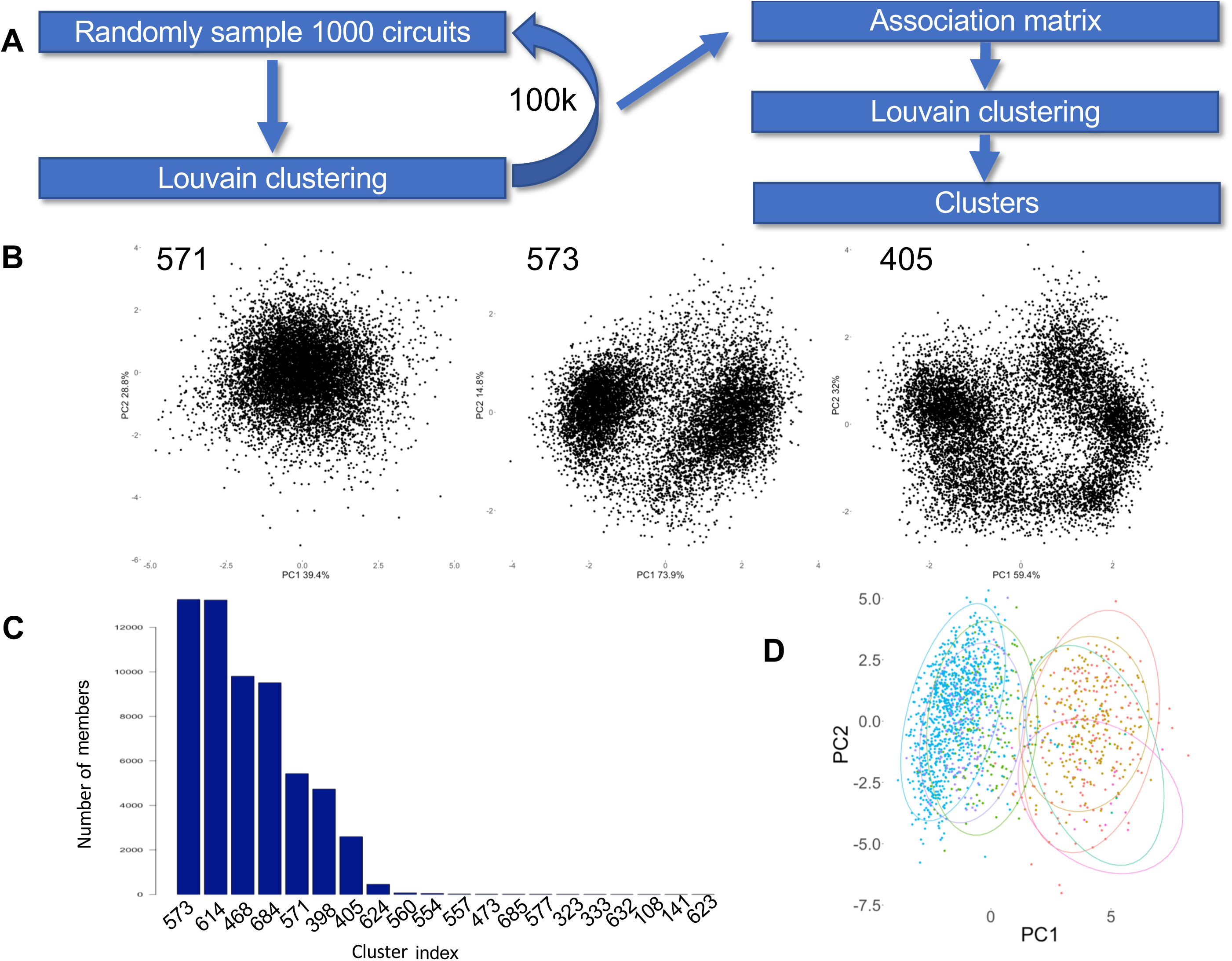
Clustering of all non-redundant four-node gene circuits by the similarity of state distributions. **(A)** Flow chart of the clustering analysis. A subset of 1000 circuits was randomly sampled for Louvain clustering. This step was repeated for 100,000 times to generate sufficient data for constructing an association matrix. The Louvain clustering method was applied again on the association matrix to obtain circuit clusters. **(B)** The center circuits (circuit diagrams in the 1^st^ row) for three clusters (leftmost: single state; middle: two states; rightmost: circular state distribution) and the corresponding state distributions from the RACIPE simulations (scatter points in the 2^nd^ row). The numbers at the top left corner are the cluster indices. **(C)** The histogram of the number of circuits in each community with more than 10 members. **(D**) Projection of the summary statistics of the most representative circuits of every cluster onto the first two principal components. The clusters are illustrated by the ellipses of different colors.

Lastly, we generated an overview of the major circuit clusters using principal component analysis (PCA). To do so, we constructed a vector of statistics summarizing the expression of each circuit (details in the Methods section). For the most representative circuits in the largest seven clusters and projected the data to the first two principal components (**Fig.4D**). Different colors and ellipses in the PCA projection illustrate the seven major circuit clusters identified from the Louvain clustering. These circuit clusters form two groups, which are well separated by the first principal axis (PC1). The circuit clusters on the left side of PC1 corresponds to the circuits capable of generating single state distributions, while the circuit clusters on the right side of PC1 corresponds to the circuits capable of generating state distributions with multiple states. Our results from the Louvain clustering seem to provide richer details of circuit behavior than those from the PCA, while the PCA results show the relationship between the identified circuit clusters.

Taken together, this top-down approach allows us to identify major classes of circuits associated with distinct state distributions and identifies multistability as the greatest difference between the four-node circuits.

### Identifying related circuits with similar state distributions

In the previous section, we had defined a KS statistics-based distance function to quantify the difference between the state distributions from two four-node circuits. This distance function also allows comparison of any two gene expression state distributions, making it possible to identify all non-redundant four-node circuits that have the closest state distributions to any other circuit’s state distribution. Two examples are illustrated in **Fig.5**. In the first case, we started with a circuit with a state distribution of a ring of six states (**Fig.5A**). We show the top five circuits with the closest state distributions, based on the described distance function (**Fig.5B**). PCA (2^nd^ row in **Fig.5B**) and UMAP (3^rd^ row in **Fig.5B**) projections show that resulting state distributions from identified circuits indeed contain similar gene expression state distributions.

**Figure 5.**
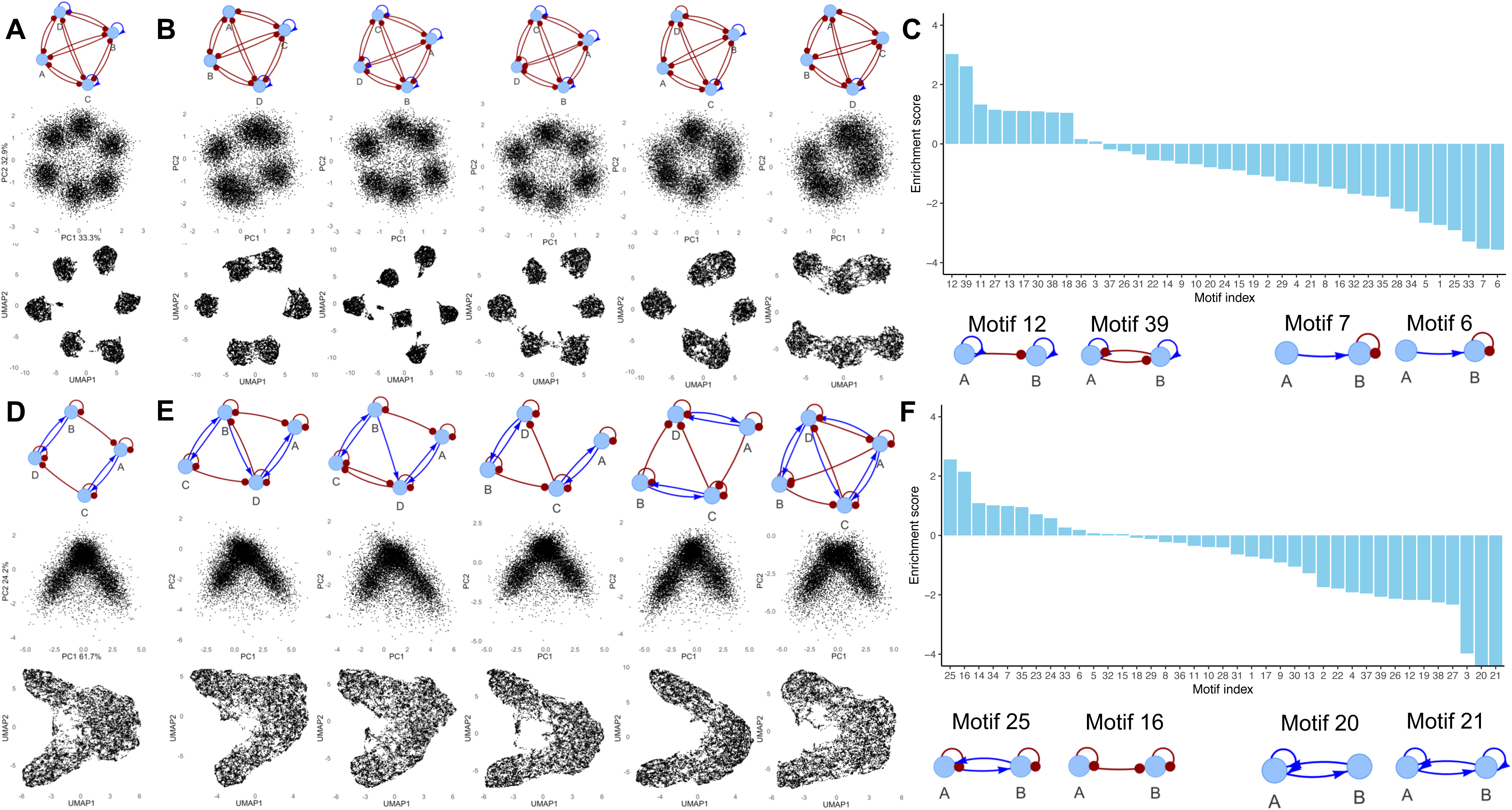
Identifying circuits with similar state distributions. A distance function based on Kolmogorov-Smirnov statistic was designed to quantify the similarity of the state distributions of two four-node gene circuits. The distance function allows us to identify other circuits with similar state distributions to a reference circuit. From these identified circuits, enrichment analysis can be applied to identify reoccurring two-node circuit motifs and their coupling. Two examples are illustrated in the plot. Panels **(A)** and **(D)** show two references circuits – **(A)** for a circuit allowing six states along a circle, and **(D)** for a circuit allow three states in a triangular shape. Panels **(B)** and **(E)** show the five most similar circuits. Each column in panels **(A)**, **(B)**, **(D)** and **(E)** shows the circuit diagram (1^st^ row), the PCA projection of RACIPE simulations (2^nd^ row), and the UMAP projection of the same data (3^rd^ row). Panels **(C)** and **(F)** show the enrichment scores of two- node circuit motifs among the topmost similar circuits (1^st^ row) and the circuit diagrams of the most over- (2^nd^ row, left side) and under- (2^nd^ row, right side) enriched motifs. Coupling results for B and E are shown in **Fig.S5** and **Fig.S6**, respectively. All enrichment results in panel **(C)** are significant (p-value < 0.05) except for motif 3, and all enrichment results for panel **(F)** are significant except for motifs 5, 8, 15, 18, 29, and 32.

We note that some of the gene-expression states may overlap in two-dimensional projections (typically with PCA), however, separation of these states usually can be discerned in other dimensions or with the projection of another method, such as UMAP. Strikingly, the identified circuits share very similar topologies – all the top five circuits have mutual inhibiting links between any two nodes and differ only by the autoregulatory links. This is in line with our previous findings that certain circuit topologies are required to generate specific structures of state distributions. Next, we performed enrichment analysis on circuits with lowest distances (top 600 similar circuits; details in Methods), from which we identified motifs 39 and 11 to be most enriched in these circuits. Here, motif 39 consists of a toggle switch with self-activation, a motif well known to generate multiple distinct states. Interestingly, motif 25, the top enriched motif for the triangularity score, is one of the least enriched motifs in this case. The results indicate that the enrichment analysis allows identification of the circuit motifs responsible for disparate state distributions. Furthermore, we observed an emphasis on positive autoregulation in the top identified motifs, which is a trend that is distinct from what was observed for earlier scores.

As a second example, we started with a circuit with a triangular state distribution of three clusters (**Fig.5D**). This state distribution was previously described by the triangularity score defined earlier (**Fig.2**). With our current analysis, we successfully identified circuits with the most similar state distributions, without using the triangularity score (**Fig.5E**). In this case, the top identified circuits, despite being very similar to one another, are structurally more different among them than those from the top six-state circuits shown (**Fig.5B**). This could be because the triangular state distributions are more commonly observed state distributions, thus more accessible to circuits of different topologies. The top three enriched motifs, as shown in **Fig.5F**, are the same motifs as identified from our earlier analysis using the triangularity score (**Fig.3**). In particular, the same motif 25 was identified once again as the topmost enriched motif for the triangular state distribution. These outcomes demonstrated the effectiveness of the distance function in identifying circuits and motifs responsible for similar state distributions. Furthermore, we have shown with two orthogonal methods that motif 25 is implicated in generating a triangular distribution of three states.

### Identifying small core regulatory circuits from single-cell gene expression

We have demonstrated how our approach can identify four-node circuits with similar state distributions to other circuit’s state distribution. Now, we extended our analysis to identify four-node circuits with similarities to experimentally observed state distributions from single-cell RNAseq (scRNA-seq) data. This is a conceptually different analyses in that (1) the state distributions from single cell data are commonly derived from many more genes (typically the most variable genes); (2) nodes in the four-node circuits do not necessarily represent individual genes, but a contribution of a group of genes due to the potential modular structure and redundancy observed in large gene networks^44, 45^. In other words, the four-node circuits in the current study represent phenomenological models of the data. We considered circuits of four nodes here to take advantage of all simulation data generated in the current study, but this approach can be readily extended to circuits of other sizes. We also revised the KS statistics-based distance functions to enable the circuit and motif analysis for single-cell gene expression data (details in Methods).

Our method was applied to a set of scRNA-seq data from 1,720 cells of human glutamatergic neuron differentiation at week 10 post-conception^46^. **Fig. 6A** shows the PCA projection of the expression of 1448 genes to the first two principal components. This dataset consists of radial glia progenitor cells progressing through intermediate neuroblast stages to differentiated neurons. From the circuit analysis we identified the topmost phenomenological four-node circuits, the top 5 of whose circuit diagrams are shown in **Fig. 6C** and the state distribution of the topmost circuit in **Fig. 6B**. The top 20 four-node circuits are shown in **Fig.S4**.

**Figure 6.**
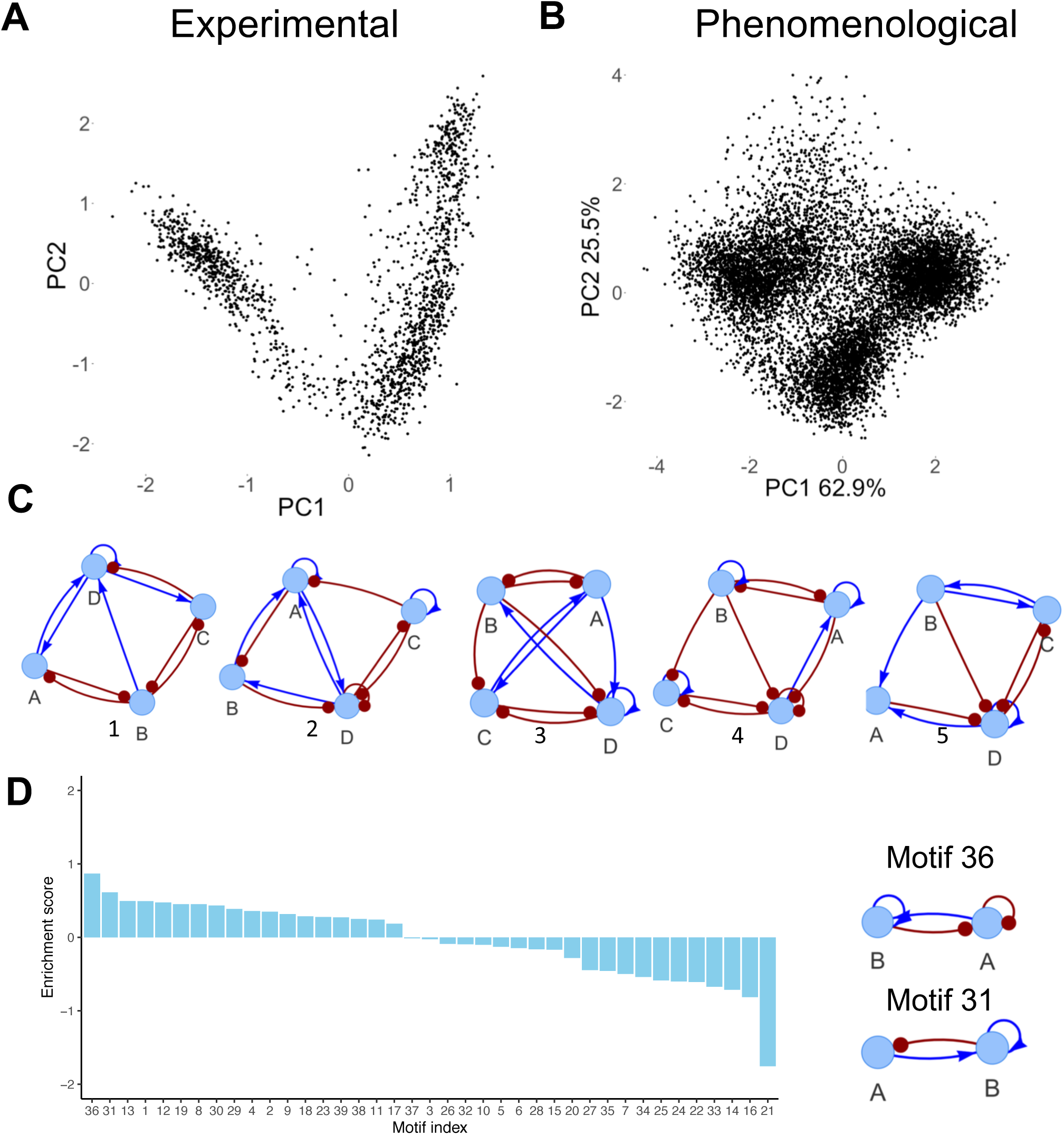
Application of the circuit analysis to scRNA-seq data of human glutamatergic neuron differentiation. **(A)** The projection of the scRNA-seq data to the first two principal components. **(B)** The projection of the simulated gene expression data of the top ranked four- node circuit to the first two principal components of the simulated data. **(C)** The diagrams of the top five ranked circuits, with ranks shown below circuits. Additional top ranked circuits are shown in **Fig.S4 (D)** The enrichment scores of two-node circuit motifs among the top-600 circuits (left panel), and the diagrams of the reoccurring two-node circuit motifs (right panel). All enrichment results are significant (p-value < 0.05) except for motifs 3, 5, 26, and 37. The motif coupling results are shown in **Fig.S7**

The circuit’s state distribution resembles that of the single-cell data in that both contain three gene expression clusters with the rightmost two clusters more connected than the leftmost cluster. These clusters potentially correspond to radial glial progenitors, intermediate progenitors, and differentiated neurons. However, the clusters from the circuit simulations appear more spherical while the experimental clusters appear more ellipsoid, presumably because of the wider range of kinetic parameters sampled in RACIPE simulations than those represented for the single cells in the experiment. From the circuit motif analysis of the top 600 circuits (**Fig. 6B**), we identified the toggle switch (motif 19) among the topmost enriched motifs. The top identified motif, motif 36, is characterized by a self-inhibiting node A and a self-activating node B, where A activates B, and B inhibits A. Upon analysis of the PCA loadings of the experimental data, we identified *VIM* as the highest negative contributor and *STMN2* and *NEUROD6* as the highest positive contributors to the first principal component. This is consistent with the experimental observation^47^ that VIM serves as a marker gene in radial glial population (leftmost cluster), and *NEUROD6* and *STMN2* are important transcription factors in intermediate progenitors and differentiated neuron populations (two clusters from the right side). We also identified *PAX6* as one of the top contributors to the radial progenitor population. *PAX6* is a TF known to play an important role in radial glial cell differentiation that activates neuronal lineages (while repressing others) to ensure correct differentiation to neurons^48^. Furthermore, it has been shown that decreasing Pax6 expression is required to turn off the neural stem-cell self-renewal program^49^. In addition, *NEUROD6* has also been shown to be implicated in sustaining the gene expression program of neurons and for promoting differentiation by triggering cell cycle withdrawal^50, 51^.

Remarkably, the top identified motif (#36) is consistent with the regulatory interactions responsible for neuronal cell differentiation^47–51^. In this circuit motif, one node (network involving *PAX6*) decreases its own expression and activates another node (network involving *NEUROD6),* which activates itself and represses the other node. Interestingly, despite the general triangular shape of the experimental state distribution, these enriched motifs identified here were not the same as those found in the previous analysis of the triangular state distributions, suggesting that the circuit and motif analysis can recognize subtle aspects of the state distribution, such as the state locality and densities. In summary, we demonstrated the circuit analysis can be applied to experimental scRNA-seq data to identify phenomenological gene circuits capable of recapitulating experimentally observed state distributions.

## Discussion

In this study, we have developed a novel computational framework to identify gene circuits, small circuit motifs, and coupling of motifs responsible for circuit properties by evaluating their gene expression state distributions. This method can be readily generalized to model other dynamical behavior of a circuit, as long as it can be quantified by a scoring function. Our method employs the first comprehensive analysis of all four-node transcriptional regulatory circuits. We have shown how the methodology can be applied to identify circuits allowing triangular or linear state distributions, from which we can further characterize the enriched motifs and motif coupling. We have also defined a KS statistics-based distance function to quantify the differences of the state distributions between two circuits. Using this distance function, we have identified major classes of circuits with distinct state distributions, circuits with similar state distributions to other circuits, and circuits that recapitulate experimental gene expression distribution from single-cell gene expression data.

Our circuit and motif analysis has the following advantages over existing methods. First, conventional approaches defined motifs as overrepresented small circuit topologies from a large biological network. The function of the identified motifs was then analyzed by mathematical modeling and/or synthetic biology analysis of a standalone circuit motif. While this approach helps to build a fundamental understanding of motifs and their importance, it falls short to discover circuit motifs for a particular function in mind. With our approach we start out by defining a desired circuit property (such as a state distribution) and then identify two-node circuit motifs enriched in all non-redundant four-node circuits with shared features. Other recent studies^6, 14^ also utilized this motif identification strategy, however the current study provides a more quantitative and generalized methodology. While we demonstrate this with a comprehensive analysis of four-node circuits identifying two-node motifs, the method can be readily adapted to identify larger circuit motifs. Therefore, our approach can alleviate the issues of existing approaches, allowing a more robust evaluation of gene circuits according to their behavior.

Second, our method utilizes RACIPE, an ensemble-based simulation approach, to evaluate circuit behavior. Compared to the earlier methods, RACIPE allows consideration of variation in kinetic parameters present in different cells. RACIPE-simulated gene expression from an ensemble of random models are usually not randomly scattered in gene expression space but form robust clusters of models. As shown in previous studies, these clusters can usually be associated with biological relevant cellular states^26, 27, 29, 30, 52^. This definition of circuit states based on gene expression distribution is more robust compared to the conventional definition based on the steady states of dynamical systems. In this way, our approach ensures a more thorough exploration of circuit behaviors and the associated circuit motifs.

Third, our method can also be applied to infer phenomenological four-node circuit models that capture the gene expression distributions of experimental single cell data. Note that the nodes in the phenomenological models may represent the collective effects of multiple regulators, instead of individual genes^53, 54^. We have shown its application to study glutamatergic neuron differentiation. We expect that this approach is invaluable to elucidate the regulatory mechanism for systems with more complex structures of cellular states.

There are a few limitations of our current approach worth investigating in the future. First, current RACIPE modeling assumes AND logics to model regulation of multiple regulators to the same target gene. But, it is well known that circuits with different types of multivariate regulation can exhibit distinct behaviors, *e.g.*, feedforward loops with AND or OR logics ^55^. An extensive analysis on this aspect would improve our understanding of the roles of logical rules in gene circuits. Second, we focused on steady-state gene expression distributions in this study, but temporal gene expression dynamics are crucial to evaluate dynamical properties of circuits^56^, such as oscillations, excitability, and chaotic dynamics. It is possible that different types of circuits exhibit similar steady-state distributions but drastically different dynamics. Investigating both the steady-state and dynamical behaviors of gene circuits could improve our understanding of gene regulatory circuit motifs. Finally, our current analysis is illustrated with circuits of only four nodes. The approach can be generalized to analyze gene circuits or networks of larger sizes. It would be interesting to discover more complex circuit motifs and patterns of motif coupling that are not observed from the analysis of four-node circuits.

## Methods

### Generation of all four-node gene circuits

To systematically evaluate the dynamical behavior of circuit motifs, we generated all non-redundant four-node gene circuits according to the following rules. First, we obtained all possible four-node circuits, where any two genes can be connected by either an activating interaction, an inhibiting interaction, or no interaction, and any gene can have either a self-activating interaction, a self-inhibiting interaction, or no autoregulation. Here, only a maximum of one regulatory interaction was considered from one gene to another; because of the directionality of gene regulation, a maximum of two regulatory interactions is possible between two genes (*i.e.*, from the first one to the second, and vice versa). This first step leads to a total of 43,046,721 circuits. Second, circuits containing *floating*, *signal* or *target* nodes were identified and excluded, as the circuits are equivalent to those with less number of nodes. Here, a floating node is defined as a node with no interaction, neither incoming nor outgoing with another node in the circuit; a signal node is defined as a node with only outgoing interactions with other nodes; a target node is defined as a node with only incoming interactions with other nodes. These definitions hold regardless of the occurrence of autoregulation. Third, for each of the remaining circuits, we constructed an adjacency matrix, with 0 representing no interaction, 1 representing activation, and 2 representing inhibition. We then computed the trace, determinant, and eigenvalues of the adjacency matrix. We considered two circuits redundant when these values are identical. We kept one circuit from all redundant circuits, which eventually leads to a total of 60,212 non-redundant four-node gene circuits for further analysis. Nonredundant four-node circuits have been analyzed in previous work ^55^, however the sign of the interactions (*i.e.*, activation and inhibition) and autoregulation were not explicitly studied, leading to a much smaller number of circuits.

### Simulation of gene circuits by RACIPE

To explore the dynamical behavior of a gene circuit, we performed *<u>ra</u>*ndom *<u>ci</u>*rcuit *<u>pe</u>*rturbation (RACIPE) ^24^ simulations using the sRACIPE R package. RACIPE takes a network as an input and generates an ensemble of ordinary differential equations (ODEs) based models, each with a unique set of randomized kinetic parameter values. RACIPE uses a wide range of parameter values that allows for the robust exploration of circuit behavior, contrary to the conventional approaches that only explore the behavior of circuits for narrow parameter ranges.

Details of the RACIPE formulism and the implementation can be found in reference^10^. Here, each circuit was simulated with sRACIPE for 10,000 models using default parameters. The simulated data were processed with log transformation and standardization for further analysis.

### Metrics to quantify circuits with a triangular or linear structure of three states

With the simulation data, we aimed to characterize the gene expression state distribution allowed by each gene circuit topology. We first performed k-means clustering (k = 3) to the simulated gene expression data, and defined a score for triangular structure, *Q*_1_, as:

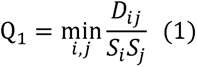

where *D_ij_* is the Euclidean distance between the centers of clusters *i* and *j*, and *S*_i_ is the average Euclidean distance of each point in the cluster *i* to the cluster center. *Q*_1_ takes the minimum of the ratio term in Equation (1) over all three pairs of clusters, *i.e.,* 1 2, 2 3, and 3 1 for *i* and *j*. When the lowest of the three ratios is still high (thus high *Q*_1_), all three clusters should be well separated. We ranked all non-redundant four-node gene circuits with *Q*_1_, so that we can identify circuits whose gene expression distribution most (or worst) resemble to a triangular structure.

Next, we defined a second score for linear structure, Q_2_, as:

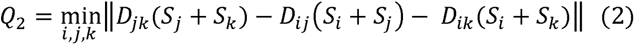

where *Q*_2_ takes the minimum of the new term over any order of the three clusters *i*, *j*, and *k*, *i.e.,* 123, 231, 312 (note that the term in *Q*_2_ is unchanged when swapping the order of *j* and *k*). The three clusters were obtained by the above-described k-means clustering. We also ranked all non-redundant four-node gene circuits with *Q*_2_, so that we can identify circuits gene expression distribution most (or worst) resemble to a linear structure.

### Circuit motif enrichment

After ranking all non-redundant four-node gene circuits with both *Q*_1_ and *Q*_2_, we explored how two-node circuit motifs are enriched in these four-node circuits from top or bottom of either ranking. To do so, we enumerated the occurrence of any two-node circuit motif (the diagrams of motifs and their indices are listed in **Fig.S1**) in all non-redundant four-node circuits. Here, for each circuit, the total number of motifs to count is 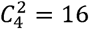. We defined an enrichment score for each circuit motif as

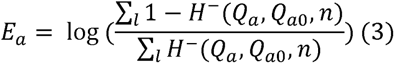

where *a* = 1 or 2 for the two scores, 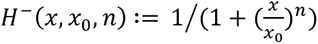 is the inhibitory Hill function, *Q_a_*_0_ is the Hill threshold, selected as the *Q_a_* value of the four-node circuit with the 600^th^ ranking by *Q_a_*. *n* is the Hill coefficient, selected as 20 to allow a sharp transition of the factor *H*^-^ from 1 to 0 for *Q_a_* near *Q_a_*_O_ . *H*^-^ is essentially a weighting factor: when *n* becomes very large, the Equation (3) becomes the log fold change of the occurrence of the circuit motif between the top 600 circuits and the rest of the circuits; a relatively small *n*, like 20, allows to consider the contributions of circuits with *Q_a_* slightly smaller than *Q_a_*_O_, to avoid the issue of zero counts. The summation from both the numerator and denominator in Equation (3) is over all non-redundant four-node circuits (,).

The significance of the enrichment was determined by a permutation test, similar to some previous approaches ^57^. A null distribution for each enrichment was created by shuffling the ranking indices of circuits for each score and applying the enrichment test. This step was repeated 10,000 times and the original enrichment results were then compared to the null distribution to estimate the p-value.

A similar approach was applied to identify enriched coupling interactions between two two-node circuit motifs over all non-redundant four-node circuits. The coupled two circuit motifs can be classified as *overlapping*, for those that share same node, and *non-overlapping*, for those that do not share same node.

### Quantifying the differences between two gene expression distributions

To quantify the differences between the RACIPE-simulated gene expression data of two four-node gene circuits (denoted as *a* and *b*), we defined a new distance function *d_ab_* as:

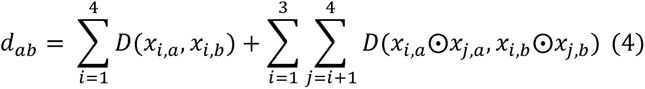

where *x_i,a_* is the expression vector of gene ; for circuit a, *x_i,a_* ⊙*x_j,a_* is the Hadamard product (element-wise product) between the expression vector of gene *i* and that of gene *j* for circuit *a*, *D*(*x,y*) denotes the Kolmogorov-Smirnov statistic^58^ between the cumulative distribution of *x* and *y*.

Furthermore, as the order of the genes in the circuits are arbitrarily assigned, an additional step was required to map the genes of the two circuits. To do so, we compute all 24 *d_ab_* where we used the default gene order for the first circuit and a permutation of gene order for the second. The lowest *d_ab_* value was eventually selected as the final distance.

### Identifying circuits with similar state distributions

Using the above-defined distance function, we constructed a matrix of pairwise distances for all the non-redundant four-node gene circuits. Starting from a circuit *a*, we can identify other circuits *b* whose *d_ab_* are the among the lowest values – these circuits are supposed to have similar state distributions.

To identify clusters of four-node circuits with similar state distributions, we adopted a subsampling approach to generate an association matrix for all the non-redundant four-node gene circuits. We performed Louvain clustering (*cluster_louvain* function in the *igraph* R package^59^) of a randomly-selected subset of 1,000 circuits for 100,000 repeats using *d_ab_* as the distance function. For every two-circuit pair, the corresponding element of the association matrix was defined as the ratio of the occurrence of the two circuits appeared in the same subset and the occurrence of the two circuits clustered together. From the association matrix we applied the Louvain clustering method again, from which we identified seven major circuit clusters with more than 500 members. For each major circuit cluster, we defined the most representative circuits as those whose state distributions have *d_ab_* of 0.05 or lower to the center circuit (circuit with lowest average KS distance to all other circuits in the community).

### Associate gene circuits with single cell gene expression data

We modified the KS test to allow for comparison of the state distributions with different number of genes. Here, we performed the KS test to compare the first two principal components of the experimental data with each two-gene combination in a four-node circuits. The distance function between a gene circuit and the experimental data was defined as the lowest distance between all the two node combinations of the synthetic circuit and the experimental data. In this way, we compared the experimental data to all the nonredundant four-node circuits, from which we identified the top ranked circuits for motif enrichment analysis.

### Data availability

Topologies of all non-redundant four-node gene circuits used in this study are available at the following link: (https://github.com/BenjaminRclauss/4node.git).

### Code Availability

R code for the RACIPE simulations, state distribution scoring, the construction of all two-node circuit motifs, and enrichment analysis of non-redundant four-node circuits is available at the following link: (https://github.com/BenjaminRclauss/4node.git)

## Acknowledgement

The study is supported by startup funds from The Jackson Laboratory and Northeastern University, and by the National Institute of General Medical Sciences of the National Institutes of Health under Award Number R35GM128717. Special thanks Christopher Baker, Gregory Carter, and Amy Yee, whom together form the Thesis Advisory committee of Benjamin Clauss.

