## Supplementary material for "A Quantitative Evaluation of Topological Motifs and Their Coupling in Gene Circuit State Distributions": SI 1

**Figure S1.** All enumerated two-node motifs and their indices.

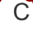

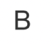

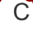

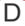

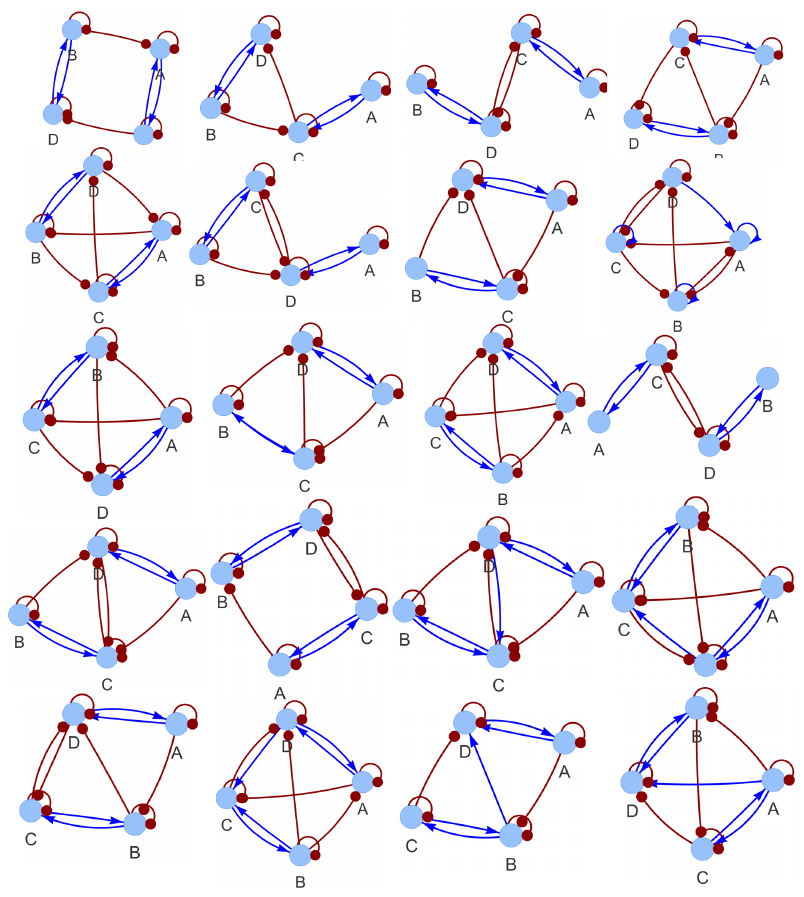

**Figure S2.** Diagrams o**f** the top 20 four-node circuits ranked highest on the triangular score from **Fig 2** arranged in decreasing order (left to right, top to bottom). Multiple examples of motif 25, occurring twice and not sharing a node are readily observed.

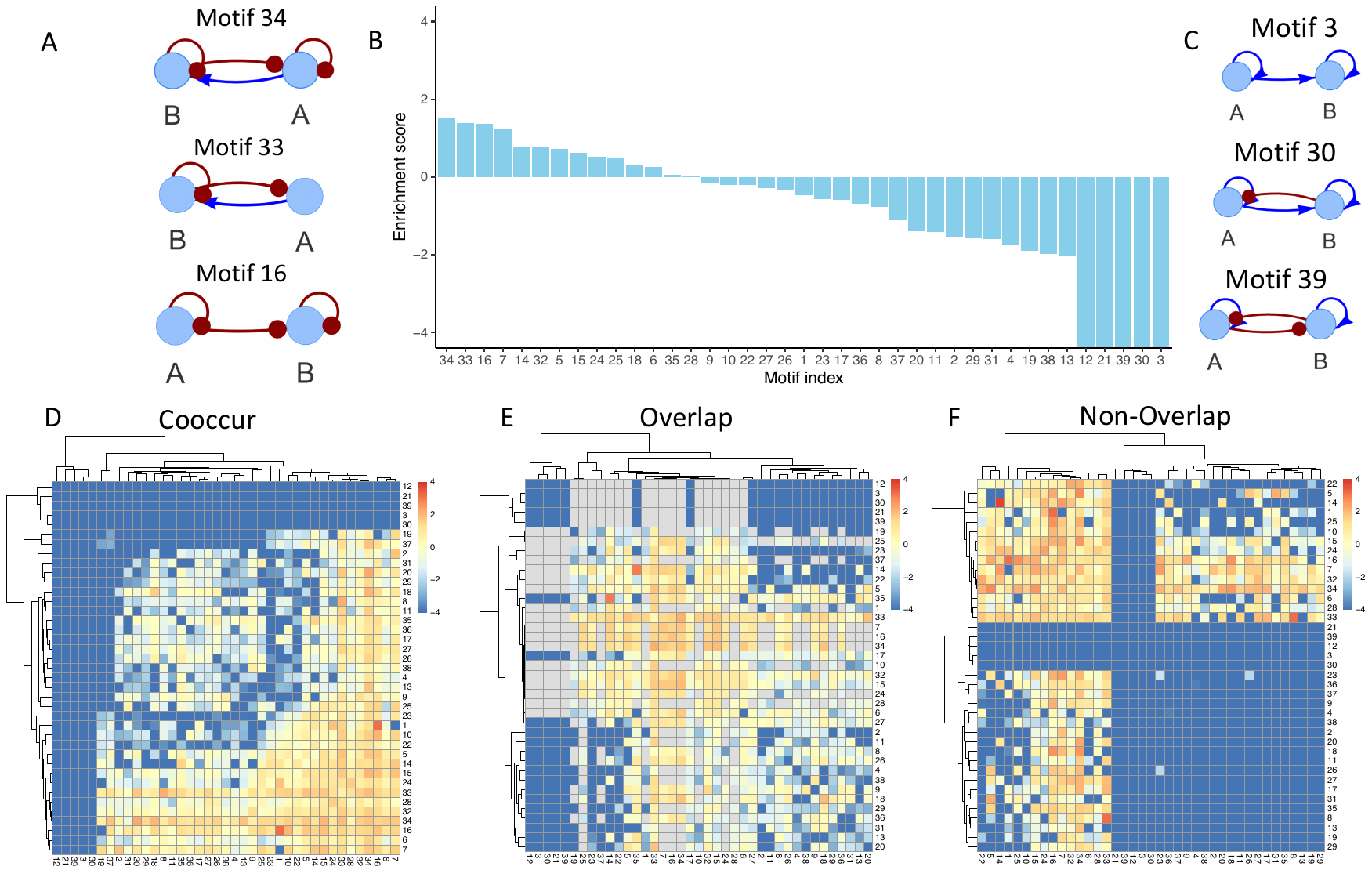

**Figure S3.** Enrichment results for the linear score comparing the top 1% of networks with the bottom 99%. **A)** Diagrams of the top three enriched two-node circuit motifs **B)** Enrichment scores for all two-node circuit motifs. Over-enriched motifs are present on the left while under-enriched motifs are present on the right. All enrichment was significant (p-value < .05) except for motifs 9, 10, 18, 28, and 35. **C)** Diagrams of the three most under-enriched motifs. Panels **D-E** are heatmaps of the enrichment scores for motif coupling. Panel **D** shows the enrichment for motifs that co-occur in the top 1% of networks; panel **E** shows the enrichment for motifs that share a node when co-occurring in the top 1%; panel **F** shows the enrichment for motifs that do not share a node when co-occurring in the top 1%.

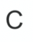

**Figure S4.** Diagrams o**f** the top 20 four-node circuits having closest state distributions of the experimental data from **Fig 6** arranged in decreasing order (left to right, top to bottom). Multiple examples of motif 36 and 31, the top two enriched two-node circuit motifs, are readily observed.

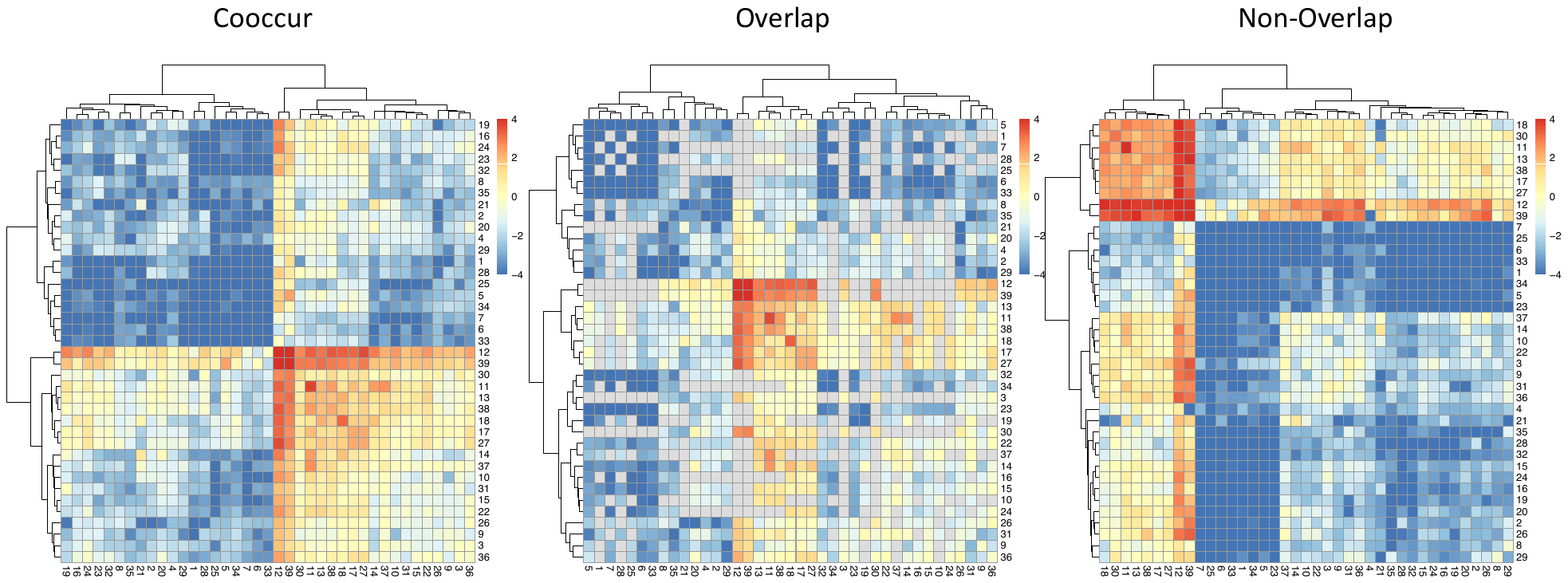

**Figure S5. Heatmaps of the enrichment scores for motif coupling for the six-state distributions, as described in Fig 5.**

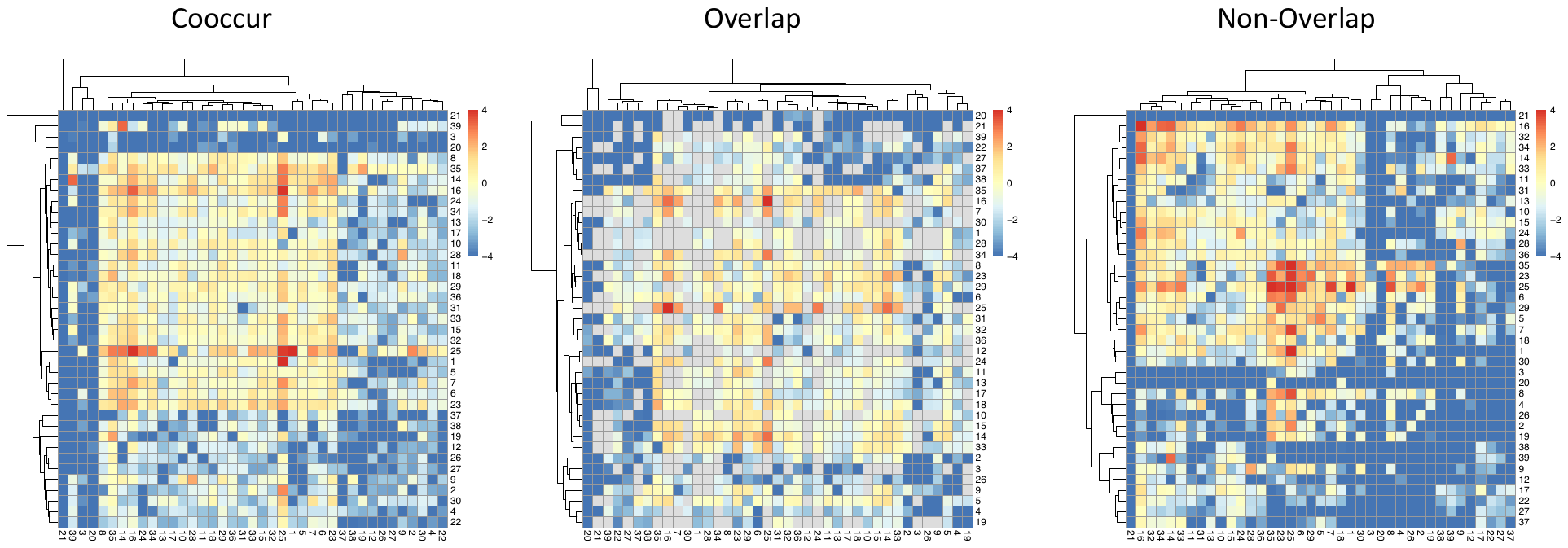

**Figure S6. Heatmaps of the enrichment scores for motif coupling for the three-state triangular state distributions, as described in Fig 5.**

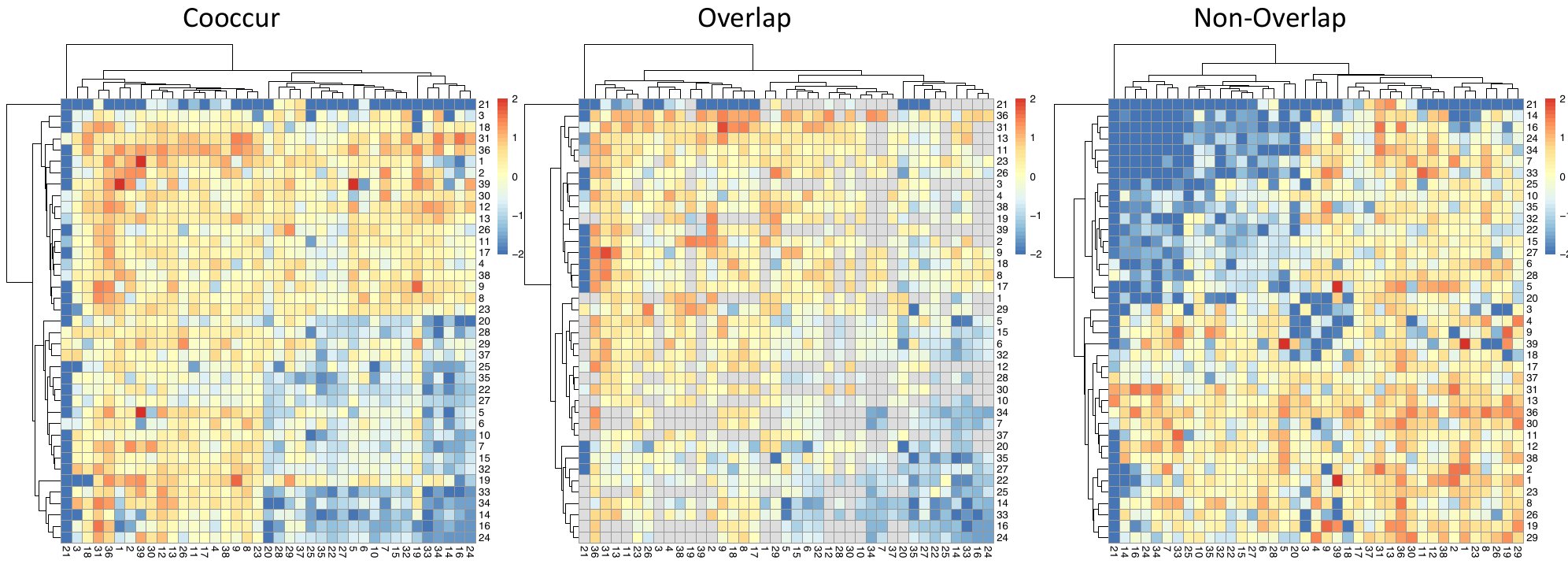

**Figure S7. Heatmaps of the enrichment scores for motif coupling for the state distributions of the experimetnal scRNA-seq data for human neuron differentiation, as described in Fig 6.**
